# Single tissue proteomics in *Caenorhabditis elegans* reveals proteins resident in intestinal lysosome-related organelles

**DOI:** 10.1101/2023.12.29.573592

**Authors:** Chieh-Hsiang Tan, Ting-Yu Wang, Heenam Park, Brett Lomenick, Tsui-Fen Chou, Paul W. Sternberg

## Abstract

The nematode intestine is the primary site for nutrient uptake and storage as well as the synthesis of biomolecules; lysosome-related organelles known as gut granules are important for many of these functions. Aspects of intestine biology are not well understood, including the export of the nutrients it imports and the molecules it synthesizes, as well as the complete functions and protein content of the gut granules. Here, we report a mass spectrometry-based proteomic analysis of the intestine of the *Caenorhabditis elegans* and of its gut granules. Overall, we identified approximately 5,000 proteins each in the intestine and the gonad and showed that most of these proteins can be detected in samples extracted from a single worm, suggesting the feasibility of individual-level genetic analysis using proteomes. Comparing proteomes and published transcriptomes of the intestine and the gonad, we identified proteins that appear to be synthesized in the intestine and then transferred to the gonad. To identify gut granule proteins, we compared the proteome of individual intestines deficient in gut granules to the wild-type. The identified gut granule proteome includes proteins known to be exclusively localized to the granules and additional putative novel gut granule proteins. We selected two of these putative gut granule proteins for validation via immunohistochemistry, and our successful confirmation of both suggests that our strategy was effective in identifying the gut granule proteome. Our results demonstrate the practicability of single tissue mass-spectrometry-based proteomic analysis in small organisms and in its utility for making discoveries.

**Significance statement:** We show that tissue-specific proteomic analysis is achievable and can be done efficiently at an individual level in a small nematode, with resolution sufficient for genetic analysis on a single animal basis. With data collected from single animals, we produced high-quality sets of proteins that described the proteomes of the gonad and the intestine. Comparison of these proteomes with the organs’ transcriptomes improved our understanding of interorgan protein transport. We applied single-tissue proteomic to describe the proteome of the gut granules in the nematode intestine, a specialized lysosome-related organelle with important functions but which is not well characterized, identifying proteins not previously known to be associated with LROs and verifying two by subcellular localization.

## Introduction

Comprising just 20 cells (1, 2), the intestine of *Caenorhabditis elegans* is not only the primary organ for nutrient uptake and storage but also the site for metal detoxification (3–7) and the synthesis and exporting of various molecules (8–10). The intestine also plays a key role in the aging process, as well as in the responses to stress and pathogens (11–13). Some of these roles depend on a distinct type of organelle commonly known as gut granules, which are characterized as a type of lysosome-related organelle (LRO) (14–17). Likely analogous to the rhabditin granules observed in other nematodes in the order Rhabditida (18–23), gut granules are robustly present in the intestinal cells (16, 23). Studies have linked them to many intestinal functions, including metabolism and storage of nutrients and trace metals, biogenesis of ascarosides, and immunity and stress-response (6, 7, 10, 24–27). However, many functions of the intestine, such as how the absorbed nutrients are passed on to other tissues and the characteristics of the gut granules, remain poorly understood.

While proteomic analyses have been applied extensively in *C. elegans* research, especially in the fields of aging (28–33) and stress response (34–36), tissue-level analyses have been hindered by the small body size of the animal. A variety of strategies have been adopted to overcome this limitation, including tissue enrichment with mutants (37, 38) and labeling proteins in specific tissues with noncanonical amino acids (39, 40) or biotin (41, 42). Analyzing tissue isolates could still offer the cleanest proteome with a much-reduced background, but it has been mostly prohibitive for somatic tissues, as they have to be manually extracted. Advancements in mass spectrometry-based proteomics have made analyses of increasingly small amounts of samples possible (43), including analyses of the *C. elegans* proteome using single worms (44–46). Individual animals are the basic units of genetic and physiological research, and the ability to perform tissue- specific proteomics at this level would be a valuable analytical tool.

We performed mass spectrometry-based proteomic analysis of manually extracted intestines and gonads of *C. elegans* and showed that most of the detected proteins can be detected using samples extracted from a single animal. We produced high-quality sets of proteins that described the proteomes of the gonad and the intestine. In a comparison of these proteomes with the organs’ transcriptomes, we identified proteins that appear to be synthesized in the intestine and then transferred to the gonad. We applied single- tissue proteomics to analyze the proteome of the gut granules by comparing the proteome of individual intestines deficient in gut granules to the wild-type. Our method identified proteins known to be exclusively localized to the granules and additional putative novel gut granule proteins, two of which- MRP-3 and W05H9.1, which we renamed LRO-1 (Lysosome-Related Organelles protein), we validate via immunohistochemistry.

Our results show that tissue-level mass-spectrometry-based proteomic analyses in small organisms, such as nematodes, can be done effectively at an individual animal level, allowing its utilization in phenotypic and genetic analyses.

## Results

### Mass spectrometry-based proteomic analysis of isolated nematode tissues

We first established a method for mass-spectrometry-based proteomic analysis of the *C. elegans* intestine and gonad using manually isolated tissues. We extracted and isolated individual intestines (Fig. 1*A* and *C*) and gonads (Fig. 1*A* and *B*) through dissection. Specifically, the intestinal tissues were severed at both ends and the gonads severed at the spermatheca to disconnect them from the rest of the worm and enable their removal. Samples were then snap-frozen and prepared “in-tube” using a “single-pot” strategy similar to those used in single-cell analysis (47). The samples were heated, sonicated, and digested with Lys-C and trypsin before mass spectrometry analysis (Fig. 1*D*, See Methods). Prior to adopting the current method, we tested other methods commonly used for preparing samples of larger quantities, all of which resulted in a very significant loss of material. The need to treat these tissues similarly to individual cells was not unexpected due to the small size of the worm and especially its organs. A single worm’s intestinal tissue contains only 20 cells, and to avoid contamination with the connected components of the alimentary system, the intestines were not extracted in their entirety.

**Figure 1.**
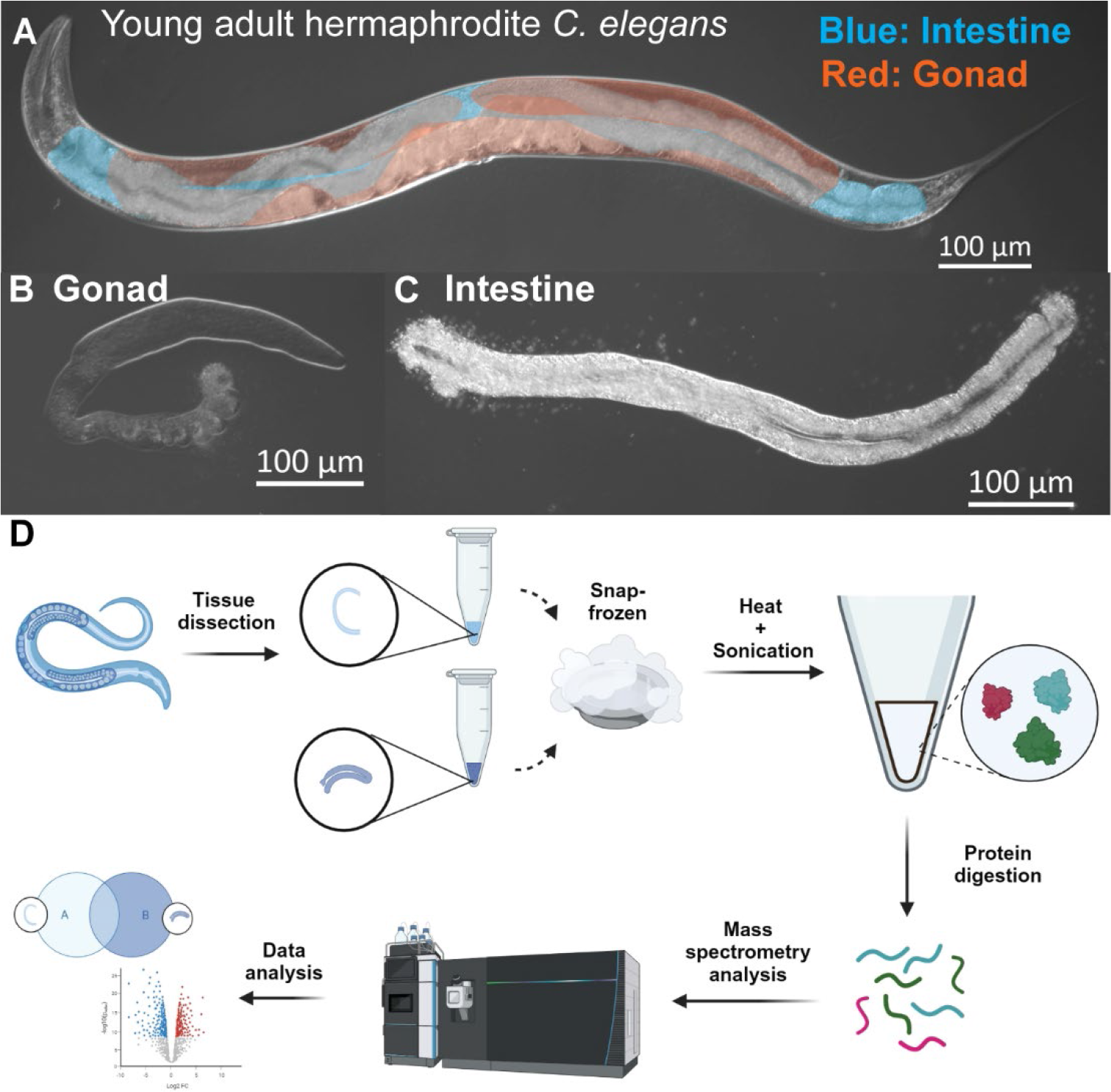
Mass-spectrometry-based proteomic analysis of extracted nematode tissues. (*A*) A young adult stage hermaphrodite *C. elegans* with the intestine (blue shade) and gonad (red shade) colored for identification. (*B*) An extracted gonad arm severed at the spermatheca. (*C*) Extracted intestinal tissue. (*D*) Schematic representation of the workflow. Intestinal tissues and gonads were extracted from young adult worms, snap- frozen, heated, sonicated, and digested prior to mass spectrometry and data analysis.

### The intestinal and gonadal proteomes of *C. elegans*

Using mass spectrometry, we detected a total of 5,962 proteins from a total of 30 mass wild-type gonad and intestine samples. Of these, 5,243 and 4,848 were identified in the gonad and the intestine, respectively (Fig. 2*A* and Dataset S1). From a single experiment with four independent samples of each organ, each consisting of tissues collected from five worms, we were able to detect a total of 5,464 proteins. Of these proteins, 4,720 and 4,267 were identified in the gonad and the intestine, respectively (Fig. 2*B* and Dataset S1). Overall, samples of the same tissue type show a high degree of similarity in both the identified proteins and their abundances while being clearly distinctive from samples of the other tissue type, albeit with notable heterogeneity across individual samples (Fig. 2*C, D* and Dataset S1).

**Figure 2.**
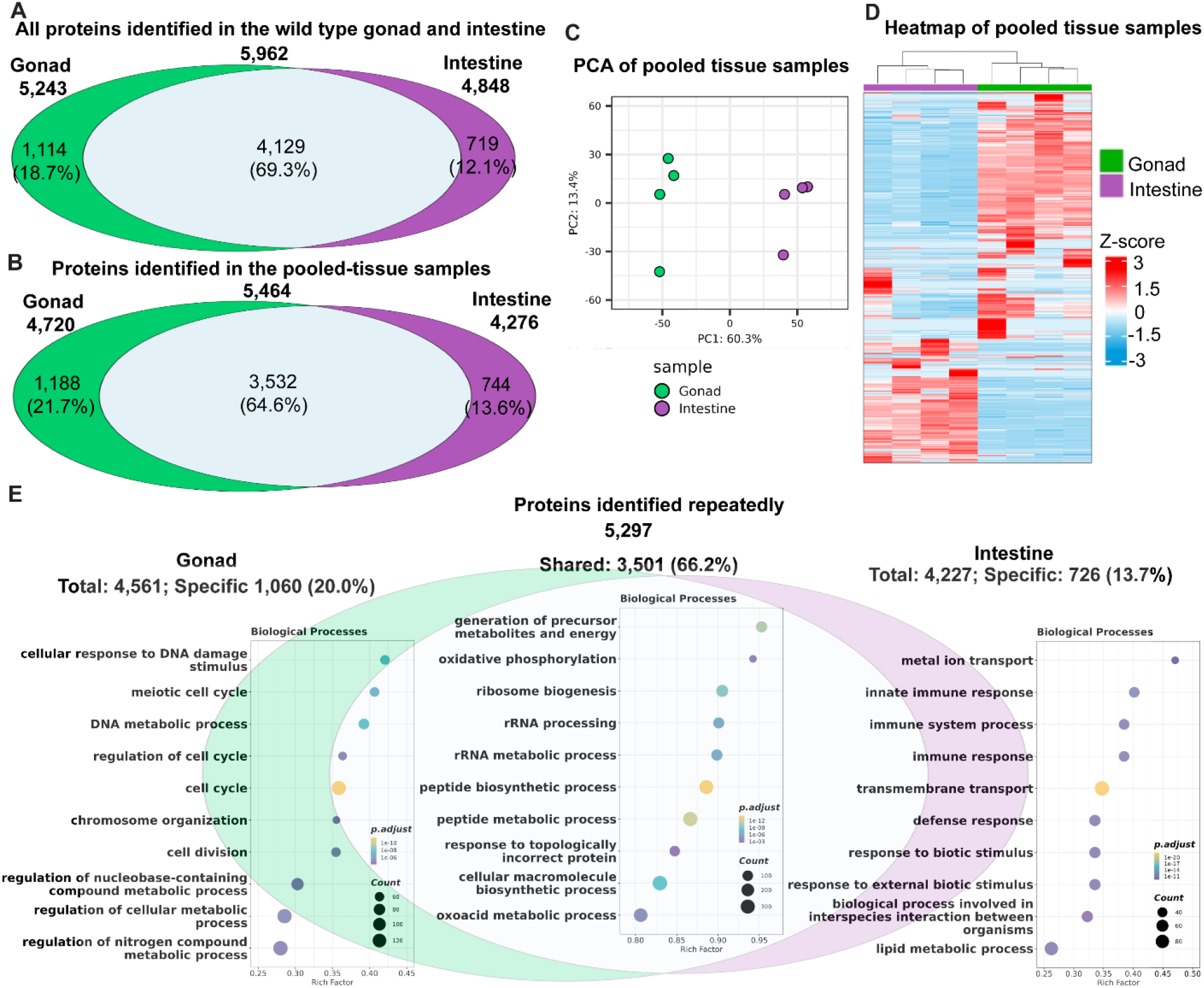
The intestinal and gonadal proteomes of *C. elegans*. (*A*) A total of 5,962 proteins were detected in our wild-type gonad and intestine samples. 5,243 were identified in the gonad, and 4,848 were identified in the intestine. 4,129 proteins were identified in both the intestine and the gonad, while 1,114 were identified only in the gonad, and 719 were identified only in the intestine. (*B-D*) Overview of proteins identified in the pooled tissue samples. Tissues were collected from five animals in each of the four samples used per tissue type. (*B*) The data from the four pooled samples for each tissue were analyzed together. From these pooled tissue samples- 5,464 proteins were detected. 3,532 of those were shared in both tissues, while 1,188 were identified only in the gonad and 744 only in the intestine. (*C*-*D*) The tissue samples of the same type show a high degree of similarity in both the identified proteins and their abundances while being clearly distinctive from those of the other tissue type. (*C*) Principal-component analysis (PCA) of the pooled tissue samples. Samples were divided into two distinct groups separated on the PC1. (*D*) Heatmap displaying the proteins of the pooled-tissue samples. The Z-score, representing the distance in standard deviations from the mean within each row, was computed by subtracting the mean protein abundance from each individual abundance and then dividing by the standard deviation across all samples in that row. (*E*) Gene Ontology biological processes enrichment analysis of proteins identified at least twice in the same tissue types. 5,297 proteins were detected more than once in the same tissue type. From these, 3,501 were shared in both tissue types, with an overrepresentation of proteins involved in essential housekeeping processes such as ribosome assembly. 1,060 were detected only in the gonad, with an overrepresentation of proteins involved in processes expected to be predominant in the germ cells, such as cell cycle. 726 were detected only in the intestine, with an overrepresentation of proteins involved in processes expected to be predominant in the intestinal cells, such as immune response, metal transport, and lipid metabolism. The Euler diagrams are area-proportional. Lists of proteins described in this figure can be found in Dataset S1.

Of the 5,962 total proteins detected, 5,297 were repeatedly detected in the same tissue type. From these, 3,501 were shared in both the intestine and the gonad, 1,060 were detected only in the gonad, and 726 were detected only in the intestine (Fig. 2*E* and Dataset S1). We subject these protein groupings to Gene Ontology enrichment analysis (48, 49), and found that, as expected, proteins involved in housekeeping processes such as ribosome assembly were enriched biological processes of the shared protein group. By contrast, proteins involved in processes expected to be prominent in the proliferating germ cells, such as cell cycle, were enriched in the group only found in the gonad. Proteins involved in processes expected to be prominent in the intestinal cells, such as lipid metabolism and immune response, were enriched in the group only found in the intestine (Fig. 2*E* and Dataset S1).

### Single-Tissue proteomic analysis

To test whether our method allows suitable coverage of the proteome, we performed mass spectrometry analysis of samples each consisting of one tissue type gathered from a single animal. We found that large portions of the tissue proteome could indeed be identified from the tissue of a single animal. Specifically, we identified a total of 4,012 proteins from six independent dissected single gonads, with up to 3,316 (82.7%) identifiable from a single gonad (Fig. 3*A* and Dataset S2). Similarly, a total of 3,732 proteins were identified from six independent dissected intestines, with up to 2,889 (77.4%) identifiable from a single intestine (Fig. 3*B* and Dataset S2). In both cases, the proteins identifiable from tissues gathered from a single worm covered over half of all the proteins we identified in our research (Figs. 2*A* and 3*A*). As with the multi-tissue samples, the tissues of the same type were much more similar to one another than they were to the other tissue both in terms of proteins identified and their abundances (Fig. 3*C, D* and Dataset S2). Similar results were also obtained from an independent trial (*SI Appendix*, Fig. S1 and Dataset S3). The proteins identified from each tissue type are also largely consistent, with nearly half of all the identified proteins identifiable in every animal, suggesting that this is a viable method for tissue proteome analysis (Fig. 3*E* and Dataset S2).

**Figure 3.**
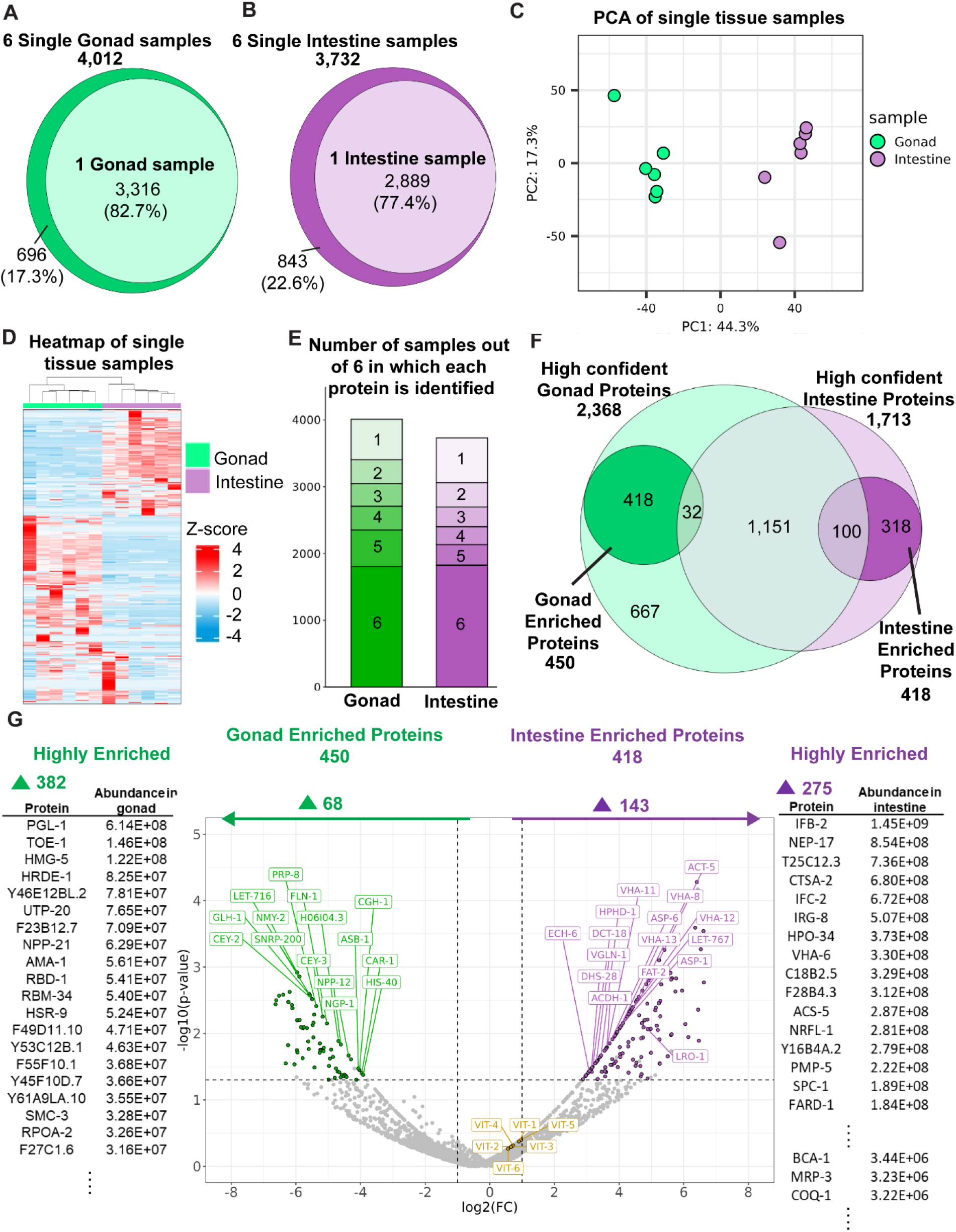
Single-tissue proteomic analysis. (*A*) A total of 4,012 proteins were identified from 6 samples of gonadal tissue, each isolated from a single animal, with up to 3,316 (82.7%) identified in a single gonad sample. (*B*) A total of 3,732 proteins were identified from 6 samples of intestinal tissue each extracted from a single animal, with up to 2,889 (77.4%) identified in a single intestinal tissue sample. (*C-D*) The single-tissue samples of the same type show a high degree of similarity and were clearly distinctive from that of the other tissue type. (*C*) Principal-component analysis (PCA) of the single tissue samples. Samples were divided into two distinct groups separated on the PC1. (*D*) Heatmap of the single tissue samples. The Z-score, representing the distance in standard deviations from the mean within each row, was computed by subtracting the mean protein abundance from each individual abundance and then dividing by the standard deviation across all samples in that row. (*E*) The vast majority of the identified proteins were repeatedly identified in tissues isolated from different animals, with nearly half of them identified in every animal. (*F*) A high-confidence set of proteins present in each tissue, 2,368 in the gonad and 1,713 in the intestine, were identified based on consistent identification of the proteins across different animals and experimental sets. Of these proteins, 450 and 418 were consistently enriched in the gonad and intestine, respectively. (*G*) Center: A volcano plot showing the relative abundance of the proteins between the gonad and the intestine. Points representing proteins in the high-confidence sets found to be enriched in the gonad (68) and intestine (143) are highlighted with the corresponding colors (green and purple, respectively), and some of the most abundant proteins are labeled. The vitellogenins, which are highly abundant in both tissues, are also labeled in yellow. Flanking the volcano plot on either side are lists of highly abundant proteins selected from those highly enriched (maxed out fold-change) in either the gonad (382) or the intestine (275), and cannot be placed in the plot. Two intestinal-enriched proteins, LRO-1 (plot) and MRP-3 (side), are also labeled and will be discussed more in later sections. The values used in this panel were based on one of the two experimental sets, but the highlighted (colored) proteins were those that were consistently enriched in the respective tissue. Green: gonad; purple: intestine; Yellow: vitellogenins. The Euler diagrams are area-proportional. Lists of proteins described in this figure can be found in Dataset S2.

Taking advantage of the significant quantities of data we collected from our single tissue mass spectrometry analysis, we compiled a “high confidence” list of proteins identified in either the gonad or the intestine (Fig. 3*F*; Dataset S2). The listed proteins were those that were consistently found in the associated tissue type (were found in at least half of datasets from each of the two trials in which each sample was a single tissue type collected from a single animal). In this way, we identified 2,368 proteins frequently found in gonad samples and 1,713 frequently found in intestinal samples. From these proteins, we then identified proteins that were consistently enriched in the gonad or the intestine (Fig. 3*G* and Dataset S2), based on the differential abundance in the two tissues (> 2-fold difference from the other tissue type in both trials). 450 and 418 proteins were found to be consistently enriched in the gonad and intestine.

### Proteome-transcriptome comparison reveals gonadal proteins with an intestinal origin

We compared our proteomic analysis to the transcriptomic data of the same tissue types by Han*, et al.* (50). Similar to our study, their sample material was also collected through dissection and physical isolation, and gene enrichment based on cross-comparison of the gonad and the intestine. Comparing the proteins that were enriched in the gonad or the intestine to the mRNAs enriched in the respective organs, we found that most of the enriched proteins (82.7% of the gonadal proteins and 62.2% of the intestinal proteins) were associated with a higher expression level of their corresponding genes in the same organ (Fig 4*A*, *B* and Dataset S4). Although parts of this difference were likely due to differences in methods, it also genuinely reflects the differences between the abundance of the proteins and the mRNA expression levels. For example, a few proteins are known to be synthesized in the intestine of *C. elegans* and then exported to other tissues. A well-established example is the vitellogenins (yolk protein precursors), encoded by six genes (*vit*-*1*∼*6*) that are expressed exclusively in the intestine (51, 52), but after synthesis in the intestine, they are transported to the maturing oocyte in the gonad (8). To further investigate intestine-to-gonad protein transfer, we looked for genes that were found by Han*, et al.* (50) to be highly expressed in the intestine compared to that of the gonad but had their encoded protein present at similar levels in both the intestine and the gonad (Fig 4*C* and Dataset S4). 99 proteins were identified with these characteristics, including all 6 of the vitellogenins. Other such proteins include: NPA-1, the *C. elegans* ortholog of the nematode polyprotein allergen/antigens proteins (NPA). Members of NPAs in parasitic nematodes bind lipids and are hypothesized to be involved in lipid distribution (53). Transthyretin-like proteins- TTR-35, 41,45,46, members of which have been suggested to be associated with the *C. elegans* yolk proteome (54). Lipid transport and storage proteins such as LBP-6 and ACBP-3; and predominantly intestinally expressed proteins with germline-associated phenotypes, such as PUD-1.2, PUD-2.2, and DCT-16 (28, 55, 56), and metabolic enzymes (Fig. 4*D* and Dataset S4).

**Figure 4.**
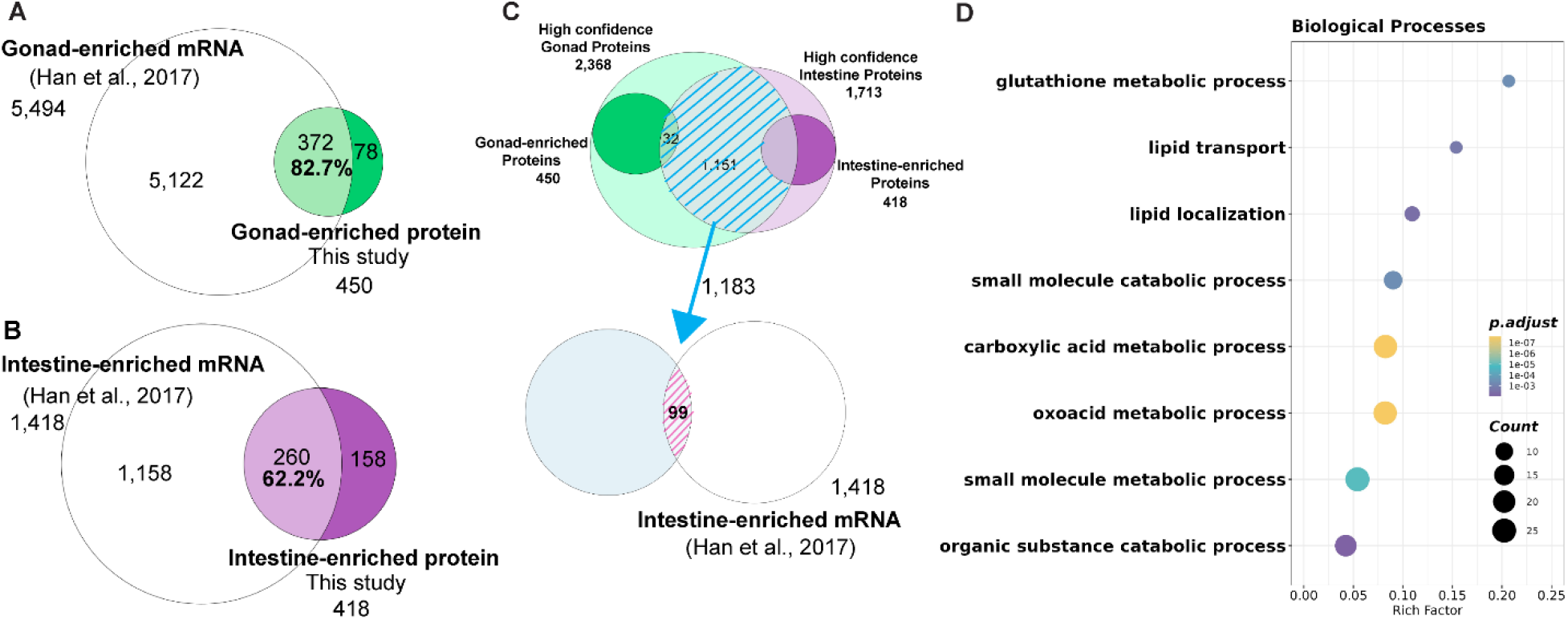
Proteome-transcriptome comparison reveals gonadal proteins with an intestinal origin. (*A*-*B*) The majority of the genes encoding the proteins enriched in the gonad (*A*) and the intestine (*B*) were also highly enriched as mRNA in the same tissues. (*C*) We identified 99 proteins that are likely gonadal proteins transcribed as mRNA in the intestine by comparing proteins that are consistently found in both the gonad and the intestine and not intestinally enriched to the list of intestinally enriched mRNAs (50) (*D*) Enrichment analysis of the list of 99 likely transported from the intestine to the gonad features proteins involved in lipid transport and localization as well as other metabolic processes. The Euler diagrams are area-proportional. The lists of proteins analyzed in this figure are in the Dataset S4.

### Identification of lysosome-related organelle associated proteins LRO-1 and MRP-3 through mass spectrometry-based genetic analysis

To identify proteins associated with lysosome-related organelles, we compared the intestinal proteomes of worms with and without gut granules. We hypothesize that the intestinal proteome of the gut-granule defective *glo-1* mutants would have decreased abundance of gut-granule specific proteins, since proteins normally associated with the organelle would be mislocalized and subjected to increased degradation. We thus analyzed the intestinal proteome of the *glo-1* mutant animals and compared it to the intestinal proteome of the wild type, by profiling intestinal tissue samples from single animals. In contrast to the overwhelming differences observed between the proteomes of the gonad and the intestine, the proteome of the *glo-1(-)* intestine was largely similar to that of the wild-type intestine, but with notably distinct elements (Fig. 5*A-C* and Dataset S5).

**Figure 5.**
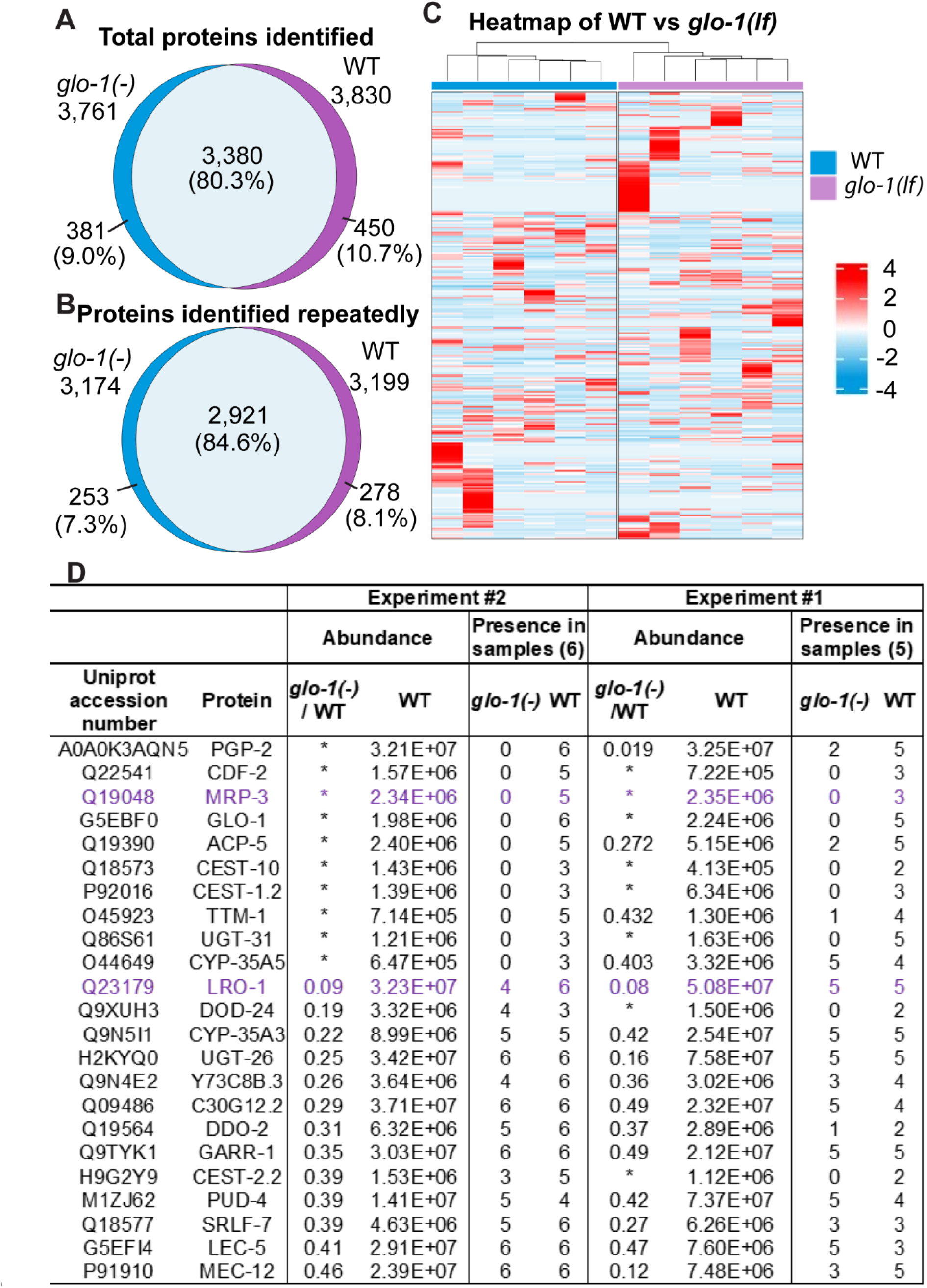
Identification of lysosome-related organelle associated proteins through mass spectrometry-based genetic analysis. (*A*) Proteins identified in *glo-1*(-) and wild- type intestines. 11 intestines of each genotype were assayed individually in two independent experiments. (*B*) Proteins repeatedly identified in the intestines of respective genotypes. (*C*) Heatmap comparing protein abundances of the *glo-1*(-) and wild-type intestines. The intestine proteome of the two genotypes is similar but distinctive. (*D*) 23 proteins, including GLO-1 were found to be consistently lower in abundance in the *glo- 1(-)* intestine than in the wild-type intestine. We hypothesized that these proteins may be part of the LRO proteome. The list was ordered on the basis of experiment 2. By the abundance of the protein in the wild-type intestines, if the protein is completely absent in *glo-1(-)* intestines and by the *glo-1(-)*/wild-type ratio otherwise. MRP-3 and LRO-1 are highlighted in purple. *: Proteins not found in the *glo-1(-)* samples. The Euler diagrams are area-proportional. Lists of proteins described in this figure can be found in Dataset S5.

To date, very few proteins were known to be gut granule specific. The cellular component term “gut granule” of Gene Ontology (48, 49) includes seven *C. elegans* proteins, of which three were only identified with the gut granules and no other cellular component. These are: PGP-2, a member of the ABC family of transmembrane transporters (24); CDF-2, a zinc transporter (6); and GLO-3, a nematode membrane protein (57). Of these three, both PGP-2 and CDF-2 have been used as markers of gut granules (26, 58, 59), and can be considered the best -characterized examples of proteins that localize to the gut granules. Comparing the proteomes found in 11 intestinal samples from individual *glo-1(-)* animals to those found in wild-type intestinal tissue samples, we identified GLO-1 and 22 other proteins that were largely missing in *glo-1* mutant tissue and so are likely part of the gut granule proteome (Fig. 5*D* and Dataset S5). The criteria were based on the differential abundance within the two genotypes (at least 50% lower in *glo-1(-)* in two independent trials) and consistent detection in the wild-type samples (at least 3 of the 6 WT samples in the 6-sample trial and at least 2 of the 5 WT samples in the 5-sample trial). Two of the 22 likely gut granule proteins we identified are PGP-2 and CDF-2, supporting our hypothesis that the *glo-1*-dependent proteins we identified are highly enriched for association with these lysosome-related organelles. Also included are three members of the carboxylesterase family, CEST-1.2, CEST-10, and CEST-2.2; CEST-2.2 is known to be involved in the synthesis of ascarosides (10), a process that involves the gut granules (60).

To further test our methodology for identifying novel lysosome-related organelles- associated proteins, we selected MRP-3, which is completely absent from the *glo-1(-)* intestine proteome, and W05H9.1, which the abundance is reduced significantly in the *glo-1(-)* intestine proteome for further validation. *mrp-3* is a member of the multi-drug resistance (MRP) subfamily of the ABC transporters superfamily (61). W05H9.1 is an uncharacterized protein highly abundant in the intestinal tissues; we renamed this protein (and gene) LRO-1 (Lysosome-related-organelle protein) based on the results of this study.

A previous biotin-based proteomic study also found LRO-1 to be enriched in the intestine and suggested that it may be localized to the cytoplasm (41). Both MRP-3 and LRO-1 are also enriched in the intestine in comparison to the gonad (Fig. 3*G*).

We modified the endogenous genomic copies of *mrp-3* and *lro-1* to label their protein products with C-terminal 3XFLAG epitope tags (62). FLAG was chosen over fluorescent proteins because of its smaller size and, thus, its reduced likelihood that appending this tag would result in mis-localization. Immunostaining was also preferable to use of fluorescent proteins due to the autofluorescent nature of the gut granules. We modified endogenous loci instead of a simpler approach of transgene expression to ensure normal germline expression (which we do not expect) and to avoid overexpression, which can cause protein mislocalization.

Immunostaining for the FLAG epitope did not label control animals and detected clear, distinct signals in the intestines but not the gonads of all of the MRP-3 and LRO-1 3XFLAG-tagged animals (Fig. 6*A*), confirming the prediction we made from our tissue-specific MS analysis of the wild-type gonad and intestine (Fig 3*G*). The staining also found both MRP-3 and LRO-1 to be subcellularly localized to granule-like organelles (Fig. 6*B*). To confirm that the observed organelles are gut granules and to validate our mass spectrometry-based predictions, we used CRISPR-Cas9 targeting to knock out the *glo-1* gene in both MRP-3 and LRO-1 3XFLAG-tagged animals. The MRP-3-3XFLAG staining was nearly completely eliminated by the knockout of *glo-1*, which validate the MS result and suggests that MRP-3 is localized nearly exclusively to the gut granules in the intestine and is likely rapidly degraded when outside of its normal environment. Similarly, the LRO-1-3XFLAG staining was greatly reduced in the *glo-1(-)* background, with significant changes in its subcellular localization. Its staining pattern shifted from granular to cytoplasmic and with decreased intensity, again validating the MS result and suggesting that LRO-1 is also a predominantly a gut granule protein.

**Figure 6.**
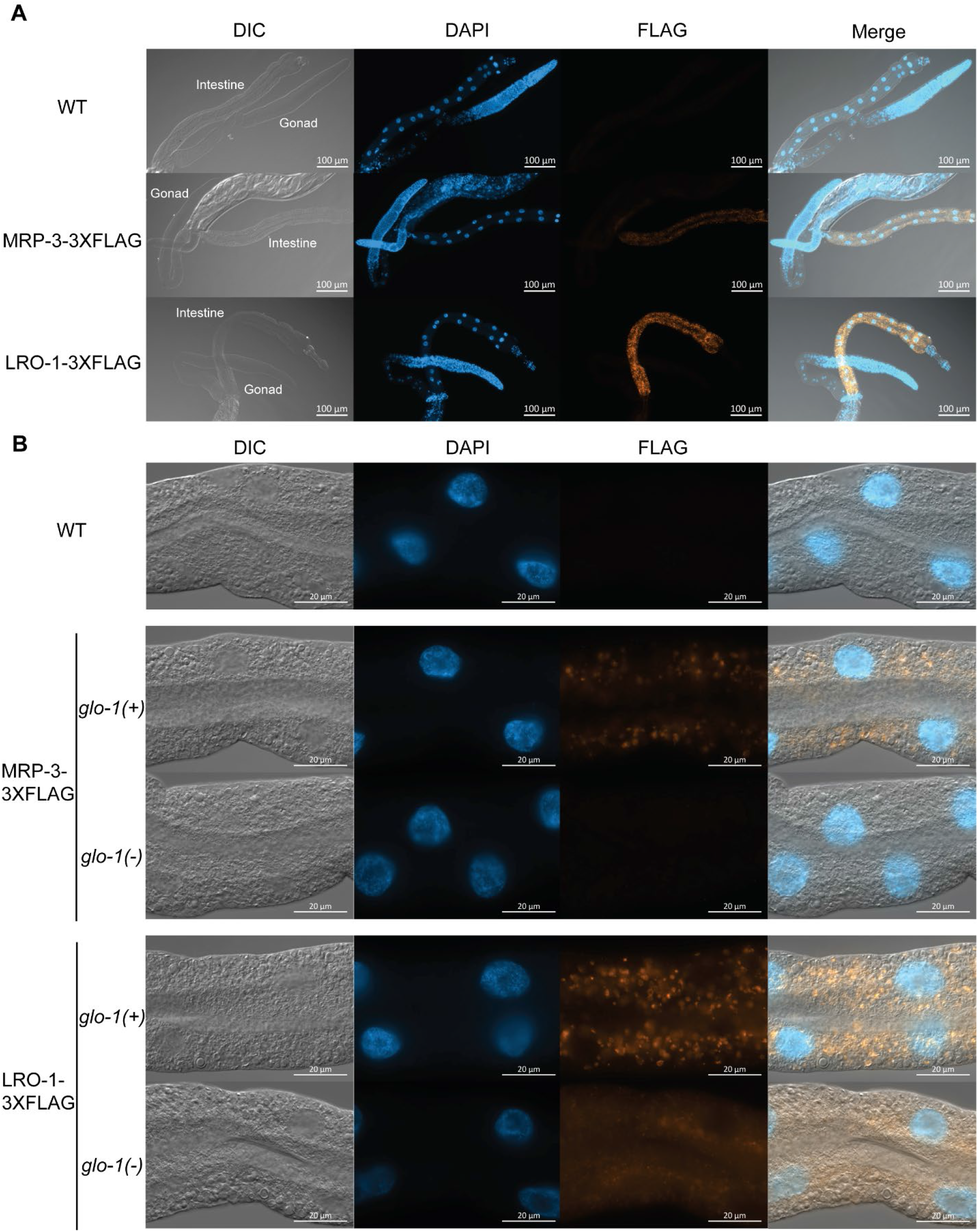
MRP-3 and LRO-1 are gut granule proteins. Anti-FLAG staining of gonad and intestine tissues dissected from wild-type control animals (WT), and from worms with a C-terminal 3XFLAG epitope inserted into the endogenous loci encoding either MRP-3 or LRO-1. (*A*-*B*) Images of the dissected tissue shown in DIC (Differential interference contrast microscopy), staining with the DNA dye DAPI (4’,6-diamidino-2-phenylindole), anti-FLAG antibody, and merged (*A*) MRP-3 and LRO-1 are abundant in the intestine but not in the gonad. Exposure was kept constant across experiments. Scale bar: 100µm (*B*) MRP-3 and LRO-1 are gut granule proteins. MRP-3 and LRO-1 are subcellularly localized to granule-like organelles in the *glo-1* (+) background; and were largely absent (MRP-3) or scattered with reduced abundance (LRO-1) in the *glo-1* (-) background, in which gut granule synthesis is impaired. Exposure was kept constant across experiments. Scale bar: 20µm

## Discussion

The small free-living soil nematode *C. elegans* is only about one millimeter in length, but it consists of a simple but clearly defined set of tissues and has for decades served as a valuable platform for a diverse array of research topics (63). As with other small organisms, however, the small size of the animal has been prohibitive for molecular analysis at the tissue level. In the case of transcriptomic analysis, while the whole animal transcriptome of *C. elegans* is well characterized and used in research early on (64–66), tissue-level transcriptomes have been a lot harder to derive. Strategies to overcome this problem include tissue enrichment with mutant alleles (67, 68), tagging or trapping mRNAs made in the target tissues (69–73), animal dissociation and FACS sorting (74–77), single-cell RNA sequencing (78, 79) as well as physically dissecting or collecting the target tissues (80–83). Lacking the amplification power provided by PCR reactions, tissue- level proteomes have been more difficult to acquire. Tissue or cell-type oriented proteomic studies in *C. elegans* have been less common but have been undertaken using strategies similar to those that have facilitated transcriptomic analysis, including tissue enrichment with mutants (37, 38) and the labeling of proteins (39–42). Using tissue isolates would offer the cleanest proteome with the lowest background, but this has been limited by the amount of samples required for proteomics. Besides one previous study of germ cells proteome, in which samples comprising over a thousand treated and untreated manually dissected gonad were analyzed, identifying a total of 839 proteins (84), tissue isolate-based analysis has only been previously performed with sperm (81) and oocytes (85), for which methods had been developed for large scale collection.

Encouraged by the recent developments in microproteomics, including proteomics analysis of single animals (44–46), we believed that tissue-level proteomic analysis of dissected tissues from *C. elegans* would now be facile, at least for larger and more coherent tissues. We focused on the intestine and the gonad, two major organs of the nematode that are relatively large and easy to isolate. Of these two, we anticipated that the intestine proteome would be easier to analyze and interpret, as the intestine is a simple organ of 20 cells that are morphologically similar to one another (1, 2). We identified approximately five thousand proteins in each of the two organs (Fig 2*A* and Dataset S1); in each organ more than four thousand proteins were identified repeatedly (Fig 2*B* and Dataset S1). We obtained relatively high coverage of each organ’s proteome, as suggested by (1) the high reproducibility of the identified proteins across samples and trials and (2) the characteristics of the proteins identified as determined via enrichment analysis (Fig 2*D* and Dataset S1). Previous whole-worm mixed-stage population studies identified up to 11,000 proteins (30, 86), so the 4000-5000 proteins we identified in a single tissue type at a single stage seem reasonable proportionally given results from transcriptomics.

We showed that mass spectrometry analyses of tissues isolated from individual *C. elegans* are feasible and provided good coverage of the proteomes targeted tissues’ proteomes sufficient to observe perturbation by mutations affecting the development of the targeted tissues (Fig. 3*A*, 3*B*, 4*A*, 4*B*, Dataset S2 and Dataset S4). We expect that this assay can be used to associate intestinal proteomics with other life characteristics, morphological or behavioral, of that individual animal.

We compiled a high-confidence list of gonadal and intestinal proteins based on data collected from individual animals and compared it to publicly available transcriptomic data. We compared these proteomes to previous transcriptome analyses to identify proteins that might be synthesized in the intestine but then transported to the gonad. The resulting set of putatively transported proteins included all six vitellogenins, which are among the very few proteins previously known to be trafficked in this manner (8), providing confirmation of this strategy. Among the 93 additional proteins identified in this manner is NPA-1, the *C. elegans* ortholog of the nematode polyprotein allergen/antigens proteins (NPA). In adult *Ascaris* nematodes, the NPA-1 ortholog is likely synthesized exclusively in the intestine (87), but is also the most abundant protein in the pseudo-coelomic fluid (88). NPA-1 and its orthologs are polyproteins containing identical or similar repeated polypeptide units, which are cleaved post-translationally (53). The derived subunits have been shown to be lipid binding and hypothesized to be involved in lipid distribution (89–93), a theory that can be supported by our data.

We utilized our method to describe the proteome of the gut granules, lysosome-related organelles known to be present in many nematodes (18–23). Gut granules have been shown to be involved in many intestinal functions. These include storage or sequestering of fat, cholesterol, heme, zinc, and copper (6, 7, 24–26); biosynthesis of ascarosides, nematode signaling molecules (10); and in immunity and stress response (27). However, many aspects of this organelle are not well understood, including its protein contents.

Based on our hypothesis that loss of LROs would lead to a decrease in the abundance of proteins that constitute its proteome, especially those with LROs as their predominant subcellular location, due to protein instability and degradation, we compared the *glo-1(-)* intestinal proteome to that of wild-type. We identified not only proteins known to be LRO proteins but also others in which the association was unknown. We selected two of the proteins for validation, MRP-3, which is undetectable in the intestines of *glo-1* mutants, and W05H9.1 (renamed LRO-1), which is greatly reduced in the intestine of *glo-1* mutants; we confirmed the LRO localization of both proteins using immunohistochemistry (Fig. 6).

In summary, in this study we describe an efficient and convenient method for the proteomic analysis of the intestinal and gonadal tissues of *C. elegans* at an individual level. With the rapid improvement of MS technologies, it is reasonable to expect other tissue types or cells of a small organism like *C. elegans* to soon be accessible for similar analysis, as long as the tissue can be physically isolated. As genomic resources improve it may also be possible to expand this work to other closely and distantly related nematodes. These other nematodes would enable the study of evolutionary divergence and could facilitate sample recovery to identify evolutionarily conserved traits that can then be experimentally probed in *C. elegans*. Promising candidate tissues for proteomic analysis include the male linker cell, a migratory cell that undergoes an atypical programmed cell death on which single-cell transcriptomic analysis has been done using dissection (83), and that of specific neurons, which have previously been shown to be accessible by dissection from the oversized nematode *Ascaris suum* (94). The intestine of *C. elegans* is arguably the core of the worms’ metabolism, homeostasis, stress, and immune responses (95). These have been exciting areas of intense research activities, including numerous transcriptomics and proteomics analyses performed at a population level, and where we envision the methods we have developed could be applied with great benefit to intestines of selected individuals displaying pertinent phenotypes.

## Materials and methods

### Nematode genetics and general methods

1. *C. elegans* were derived from the wild-type reference strain N2 (Bristol) and maintained similar to what was described by Brenner (96). Briefly, worms were cultured at 20°C on Petri plates containing Nematode Growth Medium (NGM) agar with a lawn of *Escherichia coli strain* OP50. The five times out-crossed *glo-1(zu391* lf) X (16) mutant strain OJ1347 (97) was a gift from Dr. Derek Sieburth of the University of Southern California. The following alleles were generated for this study: LGX: *glo-1(sy2102* lf), *glo-1(sy2109* lf), *lro-1(sy2059[lro-1::3xFlag])*, *mrp-3(sy2096[mrp-3::3xFlag])*.

### Intestinal and gonadal tissue isolation and preparation for mass-spectrometry (MS) analysis

One-day-old N2 (wild-type) or OJ1347 (*glo-1 (-)*) hermaphrodites were transferred on to NGM-Petri plates that did not contain a bacteria lawn, allowed to move on the surface of the agar and shed bacteria from their surface, and then transferred into a watchglass (Carolina Biological Supply Company) containing either DPBS (Dulbecco’s Phosphate Buffered Saline, modified, without calcium, magnesium, SH30378.02, Cytiva) with 0.2% DDM (n-Dodecyl-β-D-Maltoside, Thermo Fisher Scientific) or 50mM TEAB (Tetraethylammonium bromide buffer, 18597, Sigma-Aldrich) with 0.2% DDM. The worms were immobilized with levamisole (final concentration 200 µM) and cut open using with a pair of 30G 5/8” needles (PrecisionGlide, BD) in a scissors motion to extract the tissues. To separate them from the rest of the body, intestinal tissues were severed at both ends, and the gonads were cut off severed at the spermatheca. Dissected tissues were then transferred to low protein binding microcentrifuge tubes (90410, Thermo Fisher Scientific) with 0.2-0.5µL of buffer accompanying the tissue. Either 1 or 5 tissues were transferred to each tube, and flash-frozen by placing the tube in liquid nitrogen. The sample preparation method for MS analysis was modified based on the previous method as described (47). Briefly, to prepare the protein lysates, 1.5-2µL of 50mM TEAB with 0.2% DDM were added to each tube and incubate at 75°C for 1 hour. After a brief centrifugation, samples were then sonicated in a water bath for 10 min prior to digestion. The lysates were digested overnight at 37°C with 1-2 µL of the Lys-C (125-05061, Wako) and trypsin mixtures (Pierce, Thermo Fisher Scientific) (10 ng/μL each), followed by acidification through the addition of 20 μL of 1% (v/v) formic acid (FA) and 2% acetonitrile (ACN). The samples were then lyophilized and resuspended in 20 µL of 2% ACN/1% FA before lyophilizing again. Lastly, peptides were reconstituted in 10 µL of 2% ACN/0.2% FA prior to LC-MS/MS analysis.

### Mass-spectrometry (MS) analysis

LC-MS/MS analysis was performed with a Vanquish Neo UHPLC system (Thermo Fisher Scientific) coupled to an Orbitrap Eclipse Tribrid mass spectrometer (Thermo Fisher Scientific). Peptides were separated on an Aurora UHPLC Column (25 cm × 75 μm, 1.6 μm C18, AUR2-25075C18A, Ion Opticks) with a flow rate of 0.35 μL/min for a total duration of 43 min and ionized at 1.6 kV in the positive ion mode. The gradient was composed of 6% solvent B (3 min), 6-25% B (20 min), 25-40% B (7 min), and 40–98% B (13 min); solvent A: 0.1% FA in water; solvent B: 80% ACN and 0.1% FA. MS1 scans were acquired at the resolution of 120,000 from 350 to 1,600 m/z, AGC target 1e6, and maximum injection time 50 ms. MS2 scans were acquired in the ion trap using rapid scan rate on precursors with 2-7 charge states and quadrupole isolation mode (isolation window: 1.2 m/z) with higher-energy collisional dissociation (HCD, 30%) activation type. Dynamic exclusion was set to 15 sec. The temperature of the ion transfer tube was 300°C, and the S-lens RF level was set to 30. The samples described in this study were run in 3 batches. The first batch includes a trial of single wild-type gonad and intestine samples (three samples each). The data derived from this batch were used in the analysis related to Fig. 2*A*, *E*, Fig. 3*F*, *G*, Fig. 4, and Fig. S1. The second batch includes a trial of single wild-type and *glo-1(-)* intestine samples (five samples each). The data derived from this batch were used in the analysis related to Fig. 5. The third batch includes samples consisting of single wild-type gonad tissues (6 samples), single wild-type intestine tissues (6 samples), single *glo-1(-)* intestine samples (6 samples), pooled wild-type gonad tissues (5 tissues pooled in each sample, 4 samples), and pooled wild-type intestine tissues (5 tissues pooled in each sample, 4 samples). The data derived from this batch were used in the analysis related to Fig. 2, 3, 4, 5.

### Data analysis

Raw data files were analyzed using Proteome Discoverer (PD) (version 3.0, Thermo Fisher Scientific) based on the CHIMERYS algorithm against *in silico* tryptic digested Uniprot *Caenorhabditis elegans* reference proteome database containing one entry per gene (UP000001940). The maximum missed cleavages was set to 2. Dynamic modifications were set to oxidation on methionine (M, +15.995 Da) and carbamidomethylation on cysteine residues (C, +57.021 Da) was set as a fixed modification. The maximum parental mass error was set to 10 ppm, and the MS2 mass tolerance was set to 0.3 Da. The false discovery threshold was set strictly to 0.01 using the Percolator Node validated by q-value. The relative abundance of parental peptides was calculated by integration of the area under the curve of the MS1 peaks using the Minora LFQ node. The RAW data have been deposited to the ProteomeXchange Consortium via the PRIDE partner repository with the dataset identifier PXD047792 (98). The 3 batches of raw data were analyzed separately, with batch 3 analyzed in 3 separate analyses. Batch 3-1 compares the single wild-type gonad tissues (6 samples) to the single wild-type intestine tissues (6 samples). Batch 3-2 consist of the pooled wild-type gonad tissues (5 tissues pooled in each sample, 4 samples) and pooled wild-type intestine tissues (5 tissues pooled in each sample, 4 samples), as well as the samples analyzed in 3-1. Only the data from the pooled samples from 3-2 were used in this study. Batch 3-3 compares the single wild-type intestine tissues (6 samples, the same ones from 3-1) to the single *glo-1(-)* intestine samples (6 samples). Protein abundances were normalized by total peptide amount, and protein abundances (Grouped) were calculated as 2 to the power of the median log2(normalized protein abundance) in the group. The protein abundance shown in Fig. 3*G* and Fig.5*D* were group abundance. Protein ratios were calculated by the Protein Abundance Based method in PD as 2 to the power of the difference between the median log2(normalized protein abundance) in group 1 and the median log2(normalized protein abundance) in group 2. All calculations of protein abundances, protein ratio, and the statistical analysis using the t-test (background-based) were performed in PD. The PD-analyzed data used in this study can be found in Dataset S6.

The stacked bar plot and volcano plot were generated using the ggplot2 package v.3.4.4 (99). The Euler diagrams were created using ggvenn package v.0.1.10 (100) or eulerr package v.7.0.0 (101). The shared and unique protein list in Euler diagram was obtained using tidyproteomics package v.1.7.0 (102). The principal components analysis (PCA) was performed using the prcomp function from package stats 4.3.1 in R and the heatmap analysis with complete hierarchical clustering on Euclidean distances was performed using the ComplexHeatmap package v.2.16.0 (103). Both were performed with the normalized protein abundance. For Uniprot entries that correspond to multiple protein/genes, we default to the gene name included in the UniProtKB FASTA header. For GO enrichment analysis, over-representation analysis (ORA) was performed to identify the biological process using the clusterProfiler R package (104). For Fig. 2*E*, the 5,297 repeatedly identified proteins were used as the background gene set; for Fig. 4*D*, all the proteins included in the high-confidence protein sets of Fig. 3*F* were used as the background gene set. For the comparison of the transcriptome and the proteome data, the datasets were matched by gene name.

## CRISPR-Cas9 based genome editing

CRISPR-Cas9 genome editing was done with the co-conversion method as described in Arribere*, et al.* (105), to insert a 3XFlag tag sequence right before the endogenous stop codons of the endogenous locus of the genes *lro-1* and *mrp-3*, and to disrupt the function of *glo-1*. All the generated alleles were confirmed with Sanger sequencing.

For FLAG epitope coding sequence insertions, 4 to 5 mismatched silent mutations were included to prevent recutting as described in Paix*, et al.* (106), and codon-optimized based on Riddle, *et al*. (107).

The following oligonucleotides were used to generate the insertions: *lro-1(sy2059[lro-1::3xFlag])*:

Guide RNA: TCCAATAACTGCTCTACCAC

Repair template: CATCAGAAAGAGCCCCACTTCTTCCACCACCAGCAACACCGATTACAGCTCTTCCACTGGCTCCTGCCAAAGAAGAACAGCAGTTTGACTACAAAGACCATGACGGTGATTATAAAGATCATGATATCGATTACAAGGATGACGATGACAAGTAAACTTTGTTCGATAACCATGTGATTGTATTATAAAATTGTTTGCAC

*mrp-3(sy2096[mrp-3::3xFlag])*: Guide RNA: CTATTCTTAAGTAGCTCTCC

Repair template: GATGCTGGTAGAATCGTGGAGGACGGAATCCCTGGTGAACTTCTCAAGAACAGAAACTCACAGTTTTATGGTCTCGCTAAATCAGCAAAAATTGTTAATGACTACAAAGACCATGACGGTGATTATAAAGATCATGATATCGATTACAAGGATGACGATGACAAGTGATCGTTGCCGTTTTTAGTTTTTGAACAAGTTTG

To knock out *glo-1* in the *lro-1(sy2059[lro-1::3xFlag])* and *mrp-3(sy2096[mrp-3::3xFlag])* animals, we used the STOP-IN method as previously described (108). Briefly, a 43-bp universal STOP-IN cassette (GGGAAGTTTGTCCAGAGCAGAGGTGACTAAGTGATAA GCTAGC) was inserted into the first exon of the *glo-1* locus. Animals carrying a *glo-1* STOP-IN lf allele have a loss of gut granule autofluorescence phenotype indistinguishable from the loss of function allele *zu391*.

The following oligonucleotides were used to generate the insertions: Guide RNA: CTATTCTTAAGTAGCTCTCC

Repair template: GATAAAATTTCCTACAAAGTGTTGGTAATTGGTGAGGGAAGTTTGTCCAGAGCAGAGGTGACTAAGTGATAAGCTAGCTCCAGGTGTCGGTAAAACATCTATTATTCGTCG

Flanking sequence of the insertion:

Left: GATAAAATTTCCTACAAAGTGTTGGTAATTGGTGA

Right: TCCAGGTGTCGGTAAAACATCTATTATTCGTCG

The two alleles of *glo-1* knock-out, *sy2102* and *sy2109*, were separately generated but with an identical sequence.

All oligonucleotides used in this study were synthesized by IDT (Integrated DNA Technologies).

### Immunostaining of the intestine and the gonad

The immunostaining of the intestine and the gonad was performed with methods adapted from Kocsisova et al., (109) and (110) with modifications. Briefly, one-day-old hermaphrodites were transferred into a watchglass containing phosphate-buffered saline (PBS). Following washes with PBS and with PBS containing 0.1% Tween-20 (PBSTw), the worms were immobilized with levamisole (final concentration 200 µM) and cut open anteriorly with a pair of 30G 5/8” needles. The tissues were then fixed in 3% paraformaldehyde (PFA) (EM Grade, Electron Microscopy Science) in PBS for 10 minutes, transferred to a glass centrifuge tube, and post-fixed with 100% methanol at - 20°C for at least 1 hour. The fixed tissues were then rehydrated and washed three times with PBSTw, and then incubated overnight at room temperature with a monoclonal mouse anti-FLAG M2 antibody (F1804, Sigma-Aldrich) diluted 1:1000 in 30% goat serum (Thermo Fisher Scientific) in PBS. Following three washes with PBSTw, the tissues were incubated with secondary antibodies (Goat anti-Mouse Alexa Fluor 555; A-21424, Invitrogen) diluted 1:400 in 30% goat serum in PBS for 4 hours at room temperature. The tissues were then again washed three times with PBSTw and mounted on 5% agarose pad with Vectashield containing 4’,6-diamidino-2-phenylindole (DAPI) (Vector Laboratories) for observations.

Strains used for immunostaining:

Fig. 6*A*: N2: wild-type | PS10257: mrp-3(sy2096[mrp-3::3xFlag]) | PS10171: lro-1(sy2059[lro-1::3xFlag]).

Fig. 6*B*: N2: Wild-type; PS10257: mrp-3(sy2096[mrp-3::3xFlag]) | PS10271: glo-1 (sy2109), mrp-3(sy2096[mrp-3::3xFlag]) | PS10171: lro-1(sy2059[lro-1::3xFlag]) | PS10263: glo-1 (sy2102), lro-1(sy2059[lro-1::3xFlag])

### Microscopy

Images (Fig. 1, 6) were acquired with a Zeiss Imager Z2 microscope equipped with an Apotome 2 and Axiocam 506 mono using Zen 2 Blue software.

### Knowledgebases

Knowledgebases WormBase (111), the Alliance of Genome Resources (112), Gene Ontology (48, 49), and Uniprot (113) were used in the design and interpretation of experiments.

## Supporting information

Dataset S1. Data associated with Fig. 2

Dataset S2. Data associated with Fig. 3

Dataset S3. Data associated with Fig. S1

Dataset S4. Data associated with Fig. 4

Data associated with Fig. 5

All analyzed proteomic data used in this study

## Acknowledgments

We thank members of the Sternberg Lab, the Chou Lab, and PEL for discussion and suggestions. We would especially like to thank Hillel Schwartz, Nicholas Markarian, and Mark Zhang for their valuable suggestions and critical reading of the manuscript. We are also grateful for the assistance provided by Jeff Jones, Shekufeh Zareian of PEL, and Shan Li of the Chou Lab in the preliminary stages of this work, and for Wilber Palma, Stephanie Nava, and Barbara Perry of the Sternberg Lab for laboratory assistance. The mutant strain OJ1347 was a gift from Dr. Derek Sieburth of the University of Southern California. The Proteome Exploration Laboratory (PEL) is supported by the Beckman

Institute at Caltech. This research was supported by the Caltech Center for Environmental Microbial Interactions, and NIH award R24OD023041 to PWS.

## Abbreviations

LRO: lysosome-related organelle
MS: mass-spectrometry
FACS: fluorescence activated cell sorting

---

Dataset S1. Data associated with Fig. 2.

Dataset S2. Data associated with Fig. 3

Dataset S3. Data associated with Fig. S1.

Dataset S4. Data associated with Fig. 4

Dataset S5. Data associated with Fig. 5

Dataset S6. All analyzed proteomic data used in this study. Including data from every trial and including the protein abundance, normalized and without normalization.

**Figure S1.**
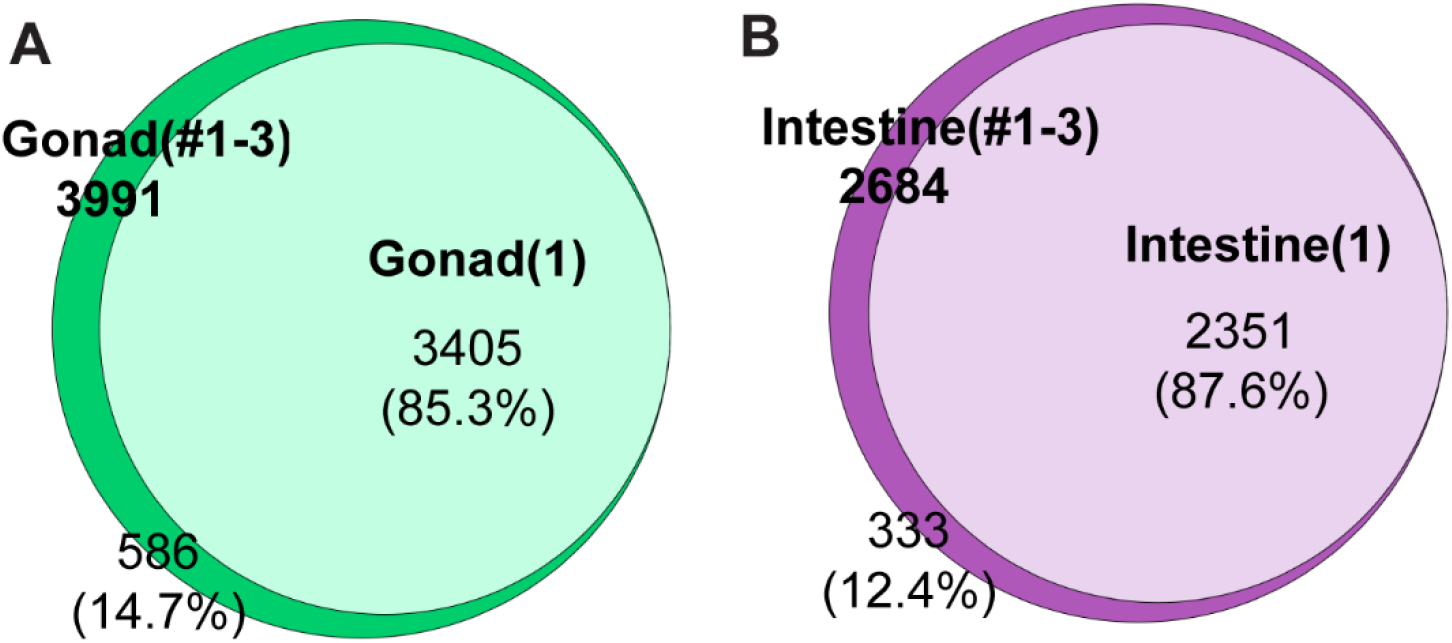
Tissue extracted from a single animal is sufficient for proteomic analysis. In an independent trial, a total of 3991 proteins were identified from 3 samples of single gonadal tissues, with up to 3405 (85.3%) identifiable in a single gonad (*A*); a total of 2684 proteins were identified from 3 samples of single intestinal tissues, with up to 2351 (87.6%) identifiable in a single intestinal tissue (*B*). The Euler diagrams are area-proportional. Lists of proteins described in this figure can be found in Dataset S3.

